# Exploring Disturbance as a Force for Good in Motor Learning

**DOI:** 10.1101/796136

**Authors:** Jack Brookes, Faisal Mushtaq, Earle Jamieson, Aaron J. Fath, Geoffrey P. Bingham, Peter Culmer, Richard M. Wilkie, Mark A. Mon-Williams

## Abstract

Disturbance forces facilitate motor learning, but theoretical explanations for this counterintuitive phenomenon are lacking. Smooth arm movements require predictions (inference) about the force-field associated with a workspace. The Free Energy Principle (FEP) suggests that such ‘active inference’ is driven by ‘surprise’. We used these insights to create a formal model that explains why disturbance helps learning. In two experiments, participants undertook a continuous tracking task where they learned how to move their arm in different directions through a novel 3D force field. We compared baseline performance before and after exposure to the novel field to quantify learning. In Experiment 1, the exposure phases (but not the baseline measures) were delivered under three different conditions: (i) robot haptic assistance; (ii) no guidance; (iii) robot haptic disturbance. The disturbance group showed the best learning as our model predicted. Experiment 2 further tested our FEP inspired model. Assistive and/or disturbance forces were applied as a function of performance (low surprise), and compared to a random error manipulation (high surprise). The random group showed the most improvement as predicted by the model. Thus, motor learning can be conceptualised as a process of entropy reduction. Short term motor strategies (e.g. global impedance) can mitigate unexpected perturbations, but continuous movements require active inference about external force-fields in order to create accurate internal models of the external world (motor learning). Our findings reconcile research on the relationship between noise, variability, and motor learning, and show that information is the currency of motor learning.

## Introduction

Neonates must determine the complex relationship between perceptual outcomes and motor signals in order to learn how to move their arms effectively. This process is repeated throughout life as humans calibrate to new environments, acquire new skills, experience neuromuscular fatigue or recover from injury. Technological advances have created robotic systems designed to accelerate the acquisition of skilled arm movements in a variety of areas including, amongst others, laparoscopic surgical training and stroke rehabilitation [1]. These devices can provide assistive forces that guide an individual’s arm through a desired trajectory or apply disturbance forces that make it more difficult for the individual to move their arm along a given trajectory.

It is now well established that providing assistive forces to neurologically intact individuals can actually impair subsequent learning [2,3]. Conversely, there is growing empirical evidence that providing disturbance forces to impair performance during training of a motor task can have a net positive effect, and lead to improved learning - enhancing performance in the task after the disturbance forces are removed [3–8]. However, formalised theoretical explanations that can account for these counterintuitive phenomena have proven elusive [9]. This is disappointing because it remains unclear how robotic devices might be best optimised in order to enhance learning (beyond this binary observation of differences between assisting and disturbing forces). The lack of a theoretical framework also makes it difficult to explain formally why assistive forces can be beneficial for individuals with neurological impairment [10], and the absence of a framework is hindering the potential utility of robotic technology in motor training. We propose that a ‘Shannon’ information theory perspective [11,12] could provide a principled approach to understanding why disruptive forces can be beneficial, and such an account could ultimately inform the development of haptic interventions.

The free energy minimization principle is the leading theoretical explanation of brain and behaviour within the domain of neuroscience, and it accounts for many empirical data within a unifying action, perception and learning framework [13–15]. The free-energy principle suggests that biological systems act to minimise free energy (an information theory measure that limits the surprise associated with sampling data). In this conceptualisation, the brain behaves as an active inference machine that formulates predictions about the environment [16]: the better the predictions about the environment, the lower the amount of free energy. Thus, the process of effective motor learning involves the system making increasingly accurate predictions about the perceptual outcome of motor commands given the current state of the system. In other words, the system will minimise entropy (the average amount of surprise) through the development of ‘forward models’ that act as neural simulators regarding how the current state of the system will respond to a given motor signal [17].

Viewed in this way, motor learning requires the system to sample information in order to extract the invariant rules that govern a range of input–output mappings [18,19]. The difficulty faced by the system relates to the large number of internal parameters that connect the sensory input to the motor output i.e. high levels of uncertainty [20]. The example of a neonate learning the mapping between perceptual and motor output illustrates how this problem can be framed from an information theory perspective. The newborn must use information generated from her exchanges with the environment in order to learn the input–output mappings and subsequently refine her predictions, so that she can successfully interact with her new surroundings. The initial reaches will be associated with high levels of uncertainty and will thus have high informational entropy (the average surprise of the outcomes sampled from the probability density). The developmental trajectory, however, will be marked by a reduction in entropy as the certainty of a predictable perceptual outcome following the generation of a motor command will increase. Thus, motor learning can be viewed as a process where entropy (i.e., uncertainty) is reduced through the development of forward models following exposure to information regarding the relationship between perceptual output and motor signal input [16].

We propose that this information perspective can account for the previous finding of superior learning outcomes from disturbance haptic force application relative to assistive guidance. Specifically, we suggest that providing assistive forces limits the amount of surprise experienced by the actor and thus constrains the amount of learning. Conversely, disturbance forces expose the individual to more information which facilitates the learning process. Following this logic, a control algorithm that provides an optimal level of surprise should lead to better learning than those that minimise uncertainty. It will be noted that a certain level of motor proficiency is required to sample information within a workspace – if an individual is unable to move their arm through the space then they will be unable to experience the surprise necessary to even start the learning process. This may explain why assistive forces have been found to help individuals with severe neurological impairment [4,21,22] or lesser skilled individuals [3,23] – as these systems allow the individual to sample the requisite information and thereby start the learning process.

Our approach is based on the idea that skilful arm movements require accurate predictions about the forces acting on the arm as it moves around the workspace. If these predictions are inaccurate then the system must contend with unexpected perturbations that will force the arm away from its desired trajectory. It has been shown that participants can learn to attenuate the impact of an unexpected perturbation in the short term by developing a ‘global impedance’ strategy, where joint stiffness rapidly increases in response to the application of a sudden unexpected force[24,25]. The development of a ‘global impedance’ strategy is a useful short term response to environments which contain unpredictable forces. Nevertheless, skilled continuous movements through a workspace require accurate forward models that allow low entropy, suggesting that the system will seek to learn (and thus predict) the underlying force field in which it is operating. On this basis, we predicted that exposure to a complex force field would, over a sufficient period, drive the system to learn how to move skilfully through the workspace (rather than adopting a short term global impedance strategy).

To test these ideas, we created a metric that quantified the information sampled as individuals learned to move their hand around an artificial environment containing a complex force field (equivalent to moving the arm through a novel viscous solution). The environment was specifically designed to produce sufficient novelty to limit the possibilities of existing forward models being adapted. These steps allowed us to examine novel motor learning in two experiments whilst providing distinct types of assistive and disturbance forces using an admittance-controlled robotic device. In our second experiment, we created a condition that would enhance learning if the Free Energy Principle inspired model has merit but would not be expected to benefit learning if the system were simply adopting a short term global impedance strategy to cope with the force field.

In our experience, there are two points worth highlighting with regard to the reported experiments. First, the experiments appear to have a similarity with a study run within Kawato’s laboratories [25]. The method section below should make it clear that the similarity is superficial. In the Kawato study, participants moved their arm along a prescribed path through a normal force field but were exposed to an unexpected perturbation when the arm diverged from the desired spatial path (resulting in participants learning to stiffen their arm in response to such perturbations). In our experiments, participants had to make continual movements through a workspace comprising a completely novel force field. This arrangement meant that our participants had to learn the underlying structure of the force field – the experiments were not about the participants moving normally and then suddenly experiencing a perturbation of an unpredictable nature. Second, our experiments included baseline measurements of how well the participants could move their arms in the novel force field. These measurements were taken before and after the participants were given the opportunity to learn the task. The baseline measures did not involve the experimental manipulations (where the robot provided assistive or disruptive forces during the learning process). Thus, the baseline measures provided an index of the motor learning that occurred throughout the experimental sessions. These measures provided the data that we needed to test the predictions of our model.

## Materials and Methods

### Participants

In Experiment 1, forty-eight right-handed participants (26 male) (M = 29.4 years, SD = 9.34 years, range 20–59 years) were recruited and randomly allocated to one of three training groups: Assistance (n = 15), Active-Control (n = 16) and Disruption (n = 17). One participant from the Active-Control group voluntarily withdrew from the experiment and their data were excluded from further analysis.

In Experiment 2, forty-six right-handed participants (25 male, aged 19 - 56 years, M = 24.93 years, SD = 6.36 years) were randomly allocated to the Adaptive Algorithm (n = 13), Adaptive Disruptive (n=17) and Random (n = 16) conditions. One participant withdrew voluntarily from the Random group after the first session and their data were not included for statistical analysis. The Psychology Research Ethics Committee at the University of Leeds approved the research.

### Procedure

In the two reported experiments using a task that required continuous tracking through a complex novel three dimensional force field, participants stood in front of a haptic robot system (HapticMASTER, see Materials) and visual stimuli were displayed on a monitor located behind the device, approximately at eye level (see Fig 1A). Two cursors were used to visually represent the actual hand and the target position of the device end-effector within the workspace on the visual display (see Fig 1D). Upon reaching the start position, the cursor started moving immediately along the first component (sub-path) for that trajectory at a constant speed of 0.1 m/s. Participants were instructed to use their preferred (right) hand to align the end-effector with a moving target as accurately as possible along pre-specified trajectories. Movement was in the Y-Z plane of the HapticMASTER system (Z – vertically upwards, Y – horizontally right relative to participant). The target cursor waited until the end of the component was reached by the participant before the next component began.

**Fig 1.**
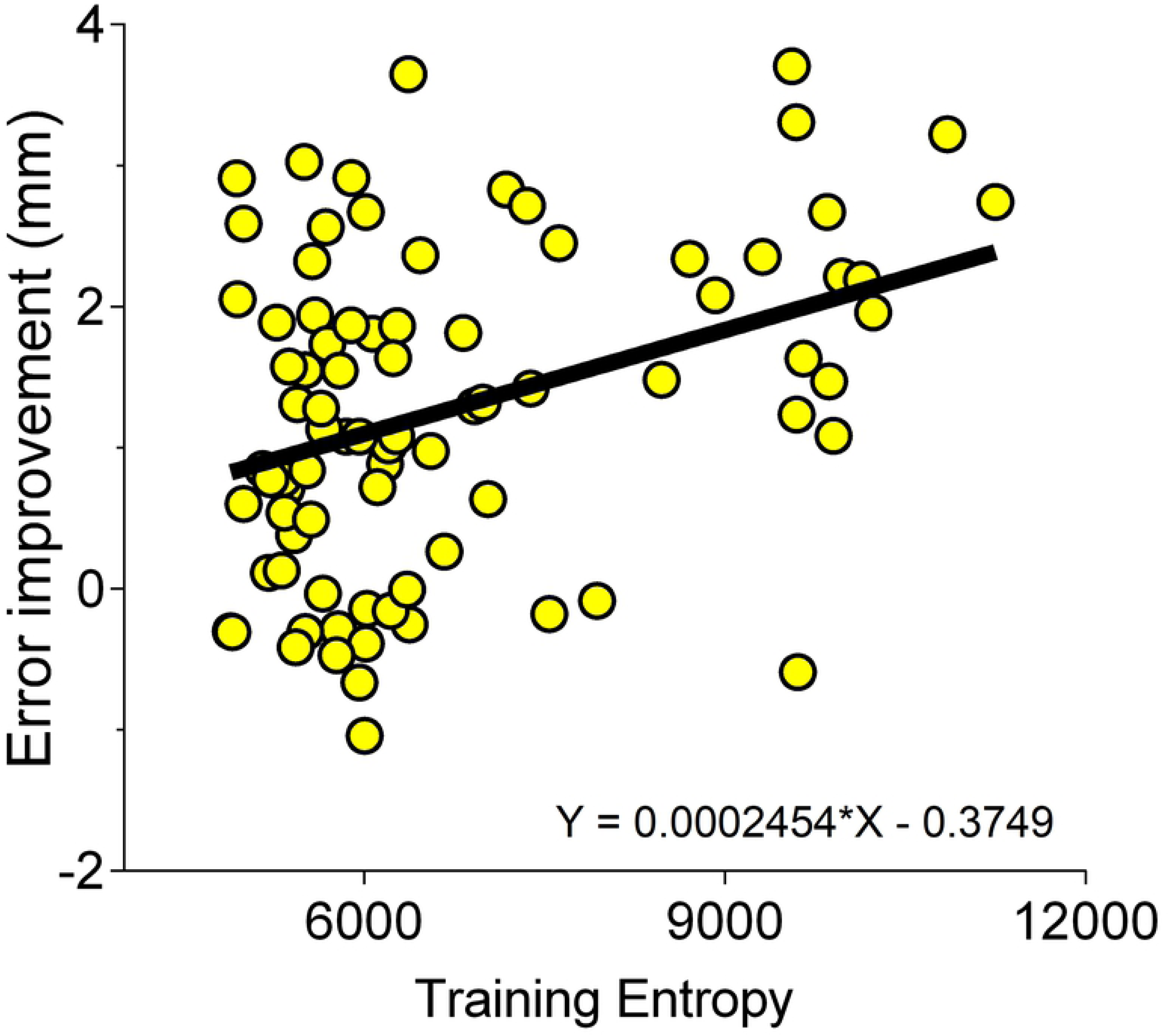
Experiment Design. (a) Plan view of the experimental setup showing the relative positions of the participant (bottom), haptic robot arm (middle) and monitor (top); (b) The target trajectories across sessions. The pre- and post-training sessions comprised 3 blocks of 10 trials following a pentagram trajectory (with no error manipulation forces). Training (across three sessions with 4 blocks of 10 trials) included error manipulation forces whilst participants navigated across a vertically rotated pentagram trajectory. (c) Quiver plot of the novel workspace force field used across all training sessions and conditions (discretized for illustrative purposes). Inset shows magnified section (approximate size 6cm x 6cm). Arrows indicate the direction and proportional magnitude of the force vector at discrete locations within the workspace. Relative magnitude is shown from white (no force) through to red (high force). (d) Blue cursor indicates the cursor (hand) position during a trial, the red circle indicates the target, the dotted black line shows the participant’s current positional error. A virtual spring sits between the cursor and the target and provides assistance, disruption, or no intervention depending on the value of *k*. N.B. Trajectory path and workspace force field remained invisible to participants throughout the experiment.

Participants were required to attend five sessions (one per day for 5 consecutive days) of approximately 15 minutes each. In sessions 1 and 5, participants followed a pentagram trajectory for three blocks of ten trials. Participants moved within a workspace force field, but had no error manipulation forces. This trajectory was based on 2D aiming tasks that have previously been used in the assessment of manual dexterity [26]. The pentagram contains five straight line components of equal length (the five edges). In Experiment 1, sessions 2 to 4 (Training) each consisted of four blocks of ten trials, with either assistive (error reducing), no or disruptive (error enhancing) forces (depending on the allocated group) superimposed over the workspace force field, following an inverted pentagram trajectory.

The target cursor was a hollow red circle, and the ‘current position’ cursor was a filled blue circle. A dotted black line was used to indicate the magnitude of the error between the current position and target cursors. To minimize fatigue, self-paced breaks with a minimum of 30 seconds rest (whilst standing or seated) were provided after each block of trials. Each session lasted approximately fifteen minutes. Experiment 2 followed same trial structure as Experiment 1, with the exception of the levels of assistance/disruption, which change trial by trial using various algorithms depending on group.

### Materials

The experiments reported here were designed to examine how error manipulation forces affect the learning of a novel workspace force field. The HapticMASTER, an admittance-controlled haptic device with a large workspace [27], was used to generate the forces and record kinematics at a rate of 1 kHz.

To simulate a novel environment, we created a workspace force field which was a function of position and calculated from the following equations:

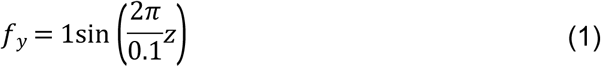

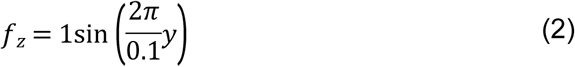

The force from the workspace force field (newtons) was a function of position (y and z, measured in meters) only. From this emerged a relatively novel environment (Fig 1C) where, in order to perform well in the task, participants needed to learn to predict the consequences of motor commands sent to the arm. Error manipulation forces (those that acted to reduce or augment execution error) were subsequently implemented using a mass-spring-damper model, as described in Equation (3):

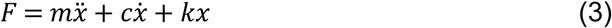

where (x) is displacement between the end effector and target positions and force is computed as a function of the distance between the actual and target positions of the end-effector. The simulation was implemented in a virtual null-gravity environment, and the end-effector mass, m, set to 3 kg and the damping, c, was set to 10 Ns/m to generate an inertial effect.

In Experiment 1, for the Active-Control condition, the stiffness k was set to 0 N/m and therefore no forces directly related to the positional error. The assistance group were provided with an assistive force implemented using *k* = 100 N/m, thereby providing full assistance, and minimizing workspace information sampling. The Disruption group had a disturbance force generated using coefficients *k* = −100 N/m, thereby providing a large prediction error for initial interactions in this condition and subsequently facilitation a larger range of movement around the workspace and information sampling.

In Experiment 2, we varied workspace information acquisition whilst also manipulating the possibility of developing a short term global impedance strategy. Specifically, we created three new training algorithms. In the Adaptive Algorithm (AA) - the virtual spring stiffness (k) varied as a function of task performance (i.e. participants had increased disturbance when performance improved and increased assistance when performance declined). The first trial of the Adaptive-Algorithm condition was always set to no intervention (*k* = 0 N/m and *c* = 0 Ns/m) in order to obtain a common benchmark measure of performance at the start of each session. The value of the stiffness coefficient at each trial was adjusted as a function of performance in previous trials, as described by Equation (4). This algorithm has been used previously as a computational model of motor adaptation to predict the force required to minimize adaptation time to a viscous environment during treadmill walking tasks [1]. In our experiment, we used the model to adjust the value of the stiffness coefficient in the current trial as a function of performance in previous trials. This allowed us to consistently keep the amount of error experienced by a participant within a small window:

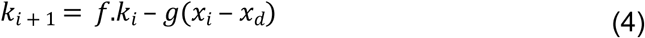

The stiffness, *k*, of the force field for the next trial is a function of the stiffness in the current trial, *i*, multiplied by a ‘forgetting factor’, *f*, and the difference between the demand error and actual error (x_d_ and x_i_, respectively), multiplied by a gain value, *g*. The values of *f* and *g* dictate the relative sensitivity of the algorithm to previous performance (captured by *k*_*i*_) and error. The sensitivity of the controller to performances obtained in previous trials is controlled by adjusting *f*: A larger forgetting factor weights the previous trials more heavily, whereas a smaller forgetting factor results in more influence from the current trial’s force field magnitude. Pilot testing informed the values of *f* and *g* to be used in the experiment and these were subsequently set at 0.5 each.

This approach allowed us to constrain the amount of information, as the level of stiffness was tuned to individual performance, constraining information by means of reducing workspace exploration since forces were always at a manageable level. The *Adaptive Disturbance* (AD) condition was identical to the AA condition, but stiffness could only decrease or stay the same between trials (i.e., the change in stiffness’ upper limit was 0). This similarly constrained information, but provided increasingly disruptive forces and therefore facilitated development of a global impedance strategy. Finally, performances in these conditions were compared against a *Random* (RAN) group - where an unpredictable stiffness value was provided (disturbance or assistance) across trials. The range of the value of k was clamped in the range −100 and 100 N/m in all 3 algorithms.

### Metrics

#### Motor Learning

Assessment (pre- and post-training) was performed without a spring stiffness (k=0), but with the same workspace force field shown in Fig 1(c). Thus, ‘learning’ can conceptually be defined as the participant’s ability to predict, and counteract, the forces arising from the workspace force field in order to minimize error. To capture how much learning occurred following training in each condition, we calculated the difference in performance in the pre- and post-training training trials. Specifically, we calculated the mean average path error scores for the three pre-test blocks and subtracted this value from the mean average path error scores from the post-test trials. Path error (E_P_) was computed as the mean Euclidian straight line distance between the end effector and the current component (sub-path) of the target trajectory. The position of the end effector was subject to a low-pass Butterworth filter (cut-off 250Hz) to remove noise in analysis of movements.

#### Analysis of Training Data

To study changes in performance as a function of training trial, we fitted a first order exponential equation to the training data using the 1^st^ order exponential fit function in the Curve Fitting Toolbox implemented in MATLAB (MathWorks Inc., Natick, MA). Training block number was used as the x value (x = 1 being the first block in the first training session), and average path error during training was used as the y value. The function uses the method of least squares to produce the most probable values of *a* and *b* in the function. The values derived from this model for each individual were subjected to group-level analysis to examine differences during training. In other words, we used the parameters of the learning function as summary statistics for random effects analysis using classical inference (i.e. ANOVA).

### Quantifying Information

To obtain a metric of information, we first parsed the workspace into discrete, independent voxels of 1 cm × 1 cm (see **Fig 2**; total size 40 cm x 40 cm). For the purposes of analysis, we created a model that assumed participants acquire information about the force output of discrete voxels, and any information acquired when the cursor was located inside a particular voxel was ‘assigned’ to that voxel. As information is accumulated for a particular voxel, newly acquired information for that voxel is discounted in value according to a weighting function. Weighting the information in this way ensures that initial “inaccurate” estimates about the expected change in force results in high amounts of surprise, and as more information is acquired, lower amounts of surprise. Effectively, the system logarithmically scales (“weights”) information in each voxel. The result of this is a metric which captures information acquired through exploration of a workspace – a higher value will result from visiting a large number of independent voxels across the workspace. The voxel size of 1cm × 1cm was a largely arbitrary selection; modelling with different voxel sizes in the range 0.25cm – 4cm shows the same pattern of results. Total weighted information gained during training can be conceptualised of as a measure of entropy.

**Figure 2.**
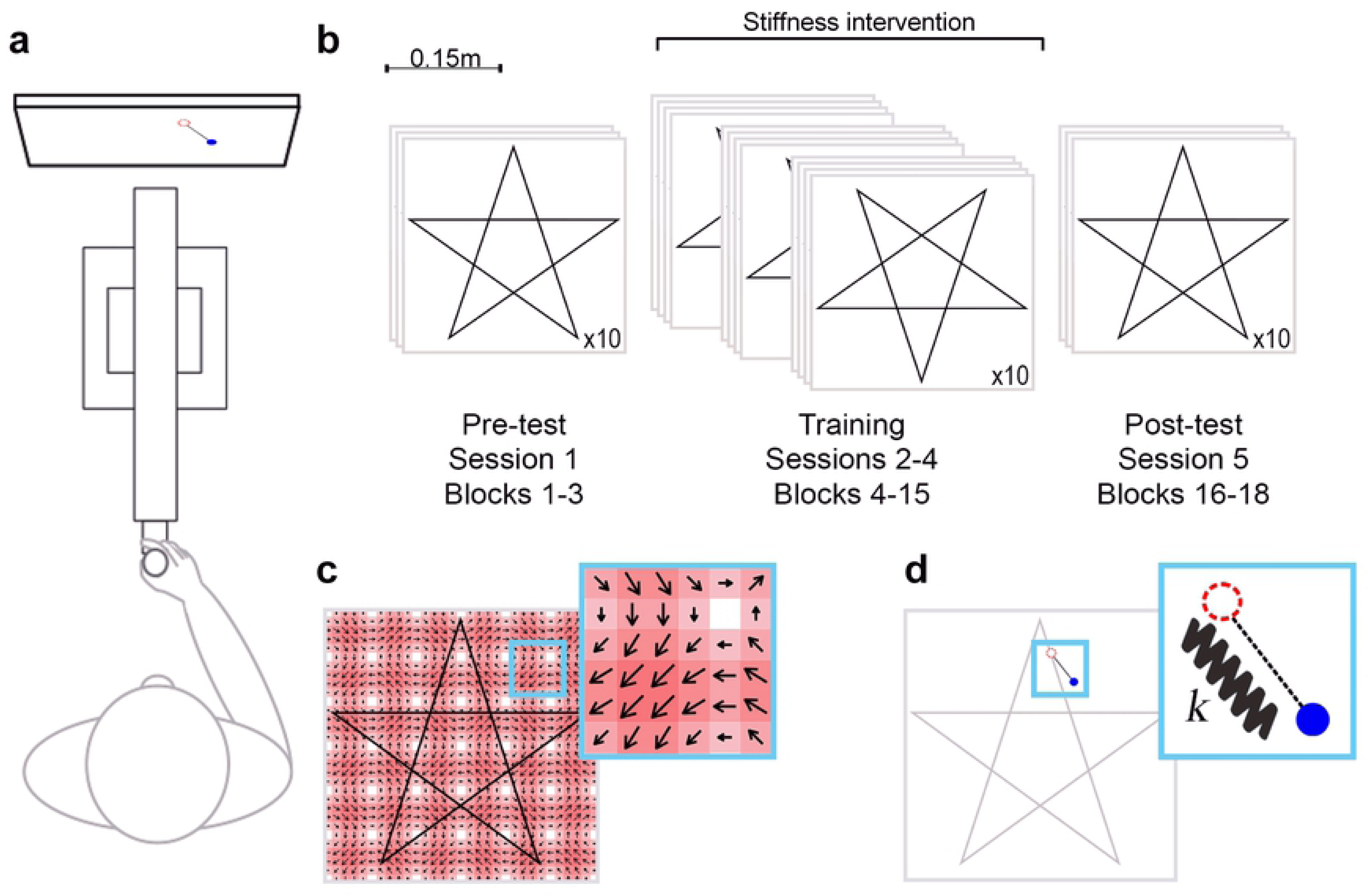
Information quantification. (a) Example simulated cursor movement across a sub-section of the workspace (10cm × 10cm). Workspace force field shown as a quiver plot, where higher force magnitude is represented by darker red shading and arrow size, and force direction indicated by arrow orientation. Workspace separated into 1cm × 1cm voxels. (b) Magnitude of change in force measured when moving along the path shown in (*a*) at a constant velocity over 1 second. Vertical black lines indicate the voxel boundary. Shaded regions under the curve separated by the vertical lines represent the information presented which is attributed to the current voxel. (c) Graphical representation of the weighting function for different values of lambda. Note that at higher values of information (in a voxel), the weighted information becomes relatively lower.

Participants were not informed about the underlying workspace force field and it remained invisible throughout the experiment. Thus, without the presence of visual information, we assumed that the sensorimotor system would have no reason to predict a change in force as a function of cursor position (at least at the outset of training). This heuristic leads to a context where the magnitude of the change in force due to the workspace force field at that point in time corresponds to a force prediction error (i.e. the difference between the experienced and predicted force). Thus, new information presented about an individual voxel was approximated as the change in force at a point in time for the voxel at the cursor position (**Fig 2b**). That is, the magnitude of change of the force vector as calculated by the workspace force field equations, Equations (**1**) and (**2**).

The information (*I*) related to a particular voxel (*i,j*) acquired throughout training up to a time *T* (total time cursor was positioned inside the voxel) was therefore:

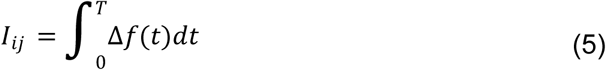

Here, information is ‘binned’ into the voxel where the end effector position is currently located (*i, j*). A value of *I* was computed for every voxel in the workspace under the assumption that information presented for a particular voxel is the magnitude of the change in force, numerically integrated over time for all points in time where the cursor position was inside that voxel (**Fig 2b**). We assumed that new information becomes less valuable as a function of the amount of information already acquired about an individual voxel as learning occurs (where models about the expected force arising from a particular voxel are updated to minimize free energy). This means that observations of changes in force have a higher probability, and therefore less surprise. Instead of using probability of sensory input estimates for each observed change in force, we opted for a more parsimonious solution by approximating surprise with a weighting function - scaling the amount of information presented to an associated information ‘value’.

The weighting method used has the desired effect for scaling information – the gradient of the weighting function = 1 when information = 0 and gradually decreases. Weighting the information in this way ensures that initial inaccurate estimates about the expected change in force results in high amounts of surprise and, as more information is acquired, the surprise is lower. The weighting formula, as a function of information presented, was:

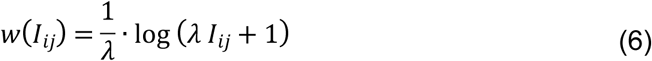

where log is the natural logarithm and *λ* corresponds to a weighting parameter. Higher values of *λ* lead to lower values of information relative to the amount of cumulative information presented, and thus faster learning about a voxel. The reported results have the value *λ* = 0.05, but we tested the model under different assumptions of *λ* (through values ranging from 0.01 to 1.00) and the pattern remained consistent.

We also assumed that the total weighted information (*TWI*) acquired was equal to the sum of the value weighted information received from each voxel of the workspace. If the workspace consists of *N*_*x*_ cells horizontally, and *N*_*y*_ cells vertically, the information value for the whole workspace at time *T* can be calculated as:

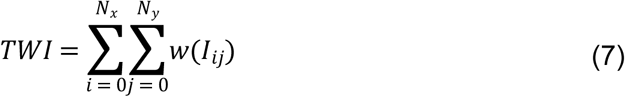

In this case the total weighted information assumes that information sampling starts at the beginning of the first training session (Session = 2) and completes at the end of the last training session (Session = 4). The total weighted information was computed per participant and is used in subsequent analyses.

It is worth noting that we could have quantified information in alternative ways to the approach described above. For example, one could model information acquisition and parameter estimation as a Kalman filter, or using Bayesian inference. However, unlike the participants in our experiments, such models would rapidly converge to the true force in a given area in only a limited number of observations. To circumvent this, we would need to make assumptions that involve including parameters estimating sensory and processing noise to slow the rate of learning. This would provide comparable results to our information scaling method if these approaches were implemented in a discrete voxel based manner (as calculated here - with exploration being rewarded as a means of sampling information and exposure to new areas of the workspace providing more information). More sophisticated models could capture the idea that repeated exposure to forces in a workspace is not sufficient for learning per se-but these also require additional assumptions e.g. an understanding (and model) of how an action (set of muscle contractions) is executed to deal with the force to maintain low positional error. Given that our aim was restricted to capturing the relationship between workspace exploration and information acquisition, we settled on a solution that provided the most parsimonious model of behavior in this task.

### Statistical Analysis

One-way between subject ANOVAs were performed to examine differences between the groups for each of the metrics described above, and Tukey’s post-hoc comparison corrected p values are reported where relevant. Partial eta squared (*η*^2^_*p*_) values are reported for effect size. We tested for, but did not find any, violations of the assumption of homogeneity of variance using Levene’s test [28]. Error bars on all Figures represent +/- 1 SEM.

## Experiment 1 – Disturbance Leads to Increased Information Sampling

We first tested the prediction that learning rates could be accelerated through the increased information provided via disturbance forces. We examined training with partially assistive (Assistance group), disturbance (Disturbance group) and no guidance (Active-Control group) forces.

In the training period, the ‘Disturbance group’ were presented with an additional force vector, whose force was generated using a negative value of *k* in the mass-spring-damper simulation. We predicted that disturbance forces would lead to (i) more surprise (as indexed by our model of information); (ii) more errors at the outset of training – indexed by *a* in the fitted function *y* = *ae*^*bx*^ and (iii) increased rate of error reduction over the training period (indexed by *b*); and finally, as a corollary of the above, (iv) superior motor learning compared (pre-post-error improvement) to the groups with lower information.

The differences in information at the early and late stages for each condition can be seen in **Fig 3**. Formal analysis of the cumulative amount of information for each group at the end of the training block revealed statistically significant differences (F (2, 44) = 34.21, p < .0001, *η*^2^_*p*_*=* .609). This effect was driven by the Disturbance group gathering more information about the workspace relative to the Active-Control (p < .0001) and Assistance (p < .0001) groups, but there was no difference between the Assistance and Active-Control groups (p = .876).

**Fig 3.**
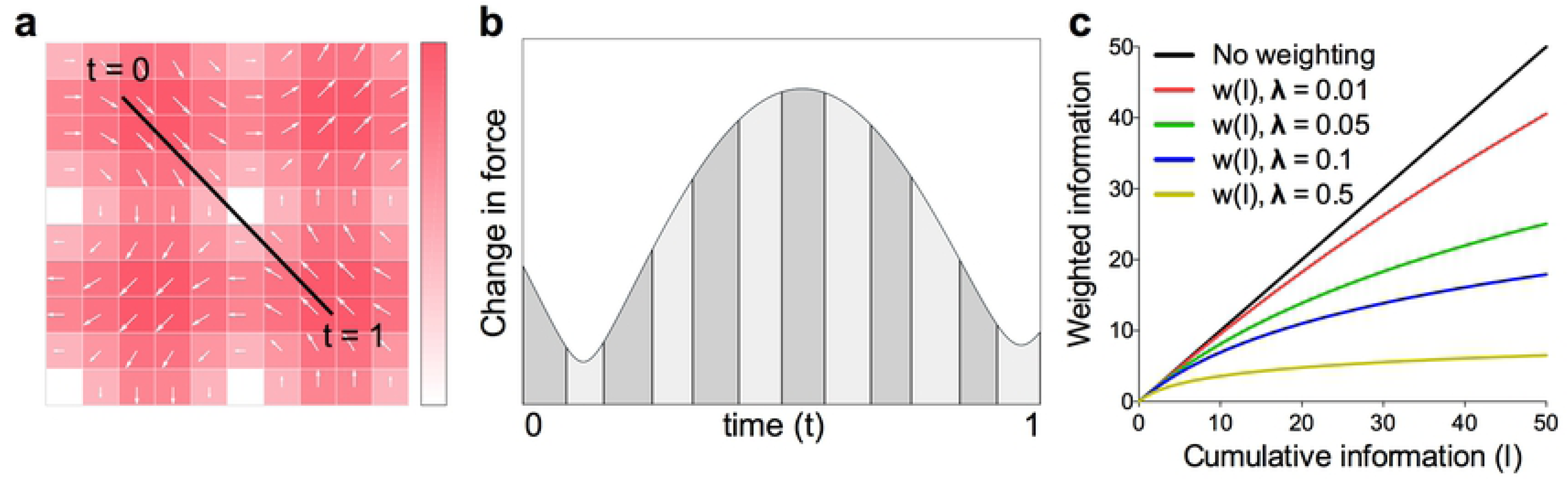
Information as a by-product of disruption. (a) The Disturbance group had more information over time at a group level; (b) Example heat maps showing the amount of information gathered across the workspace at the outset and end of training for randomly selected individual participants.

We next performed an ANOVA on the values for the exponential fit to examine differences at the outset of training. The ANOVA revealed group differences (F (2, 44) = 7.623, p = .0014, *η*^2^_*p*_*=* .257), with the Disturbance group performing worse than the Assistance group (p = .0009), although following correction for multiple comparisons, this was not significantly different to the Active-Control group (p = .1162). When comparing performance across training trials (F (2, 44) = 26.37, p <.0001, *η*^2^_*p*_*=* .545), we found that the disturbance group showed a steeper decay in error in comparison to the Active-Control (p < .0001) and Assistance Groups (p <.0001). There was no difference between learning for the Assistance and Active-Control conditions (p = .2589).

The amount of motor learning was quantified as the error improvement between the mean pre- and post-path error score (both of which were performed without any stiffness intervention [k = 0] and with the upright pentagram shape). We found significant differences in the amount of motor learning between groups (F (2,44) = 5.655, p = .0065, *η*^2^_*p*_*=* .204). Specifically, the group exposed to Disturbance forces during training on the inverted pentagram trajectory had improved significantly more than the Assistance (p = .0136) and the Active-Control (p = .0202) groups (**Fig 4**). These results are consistent with our model.

**Fig 4.**
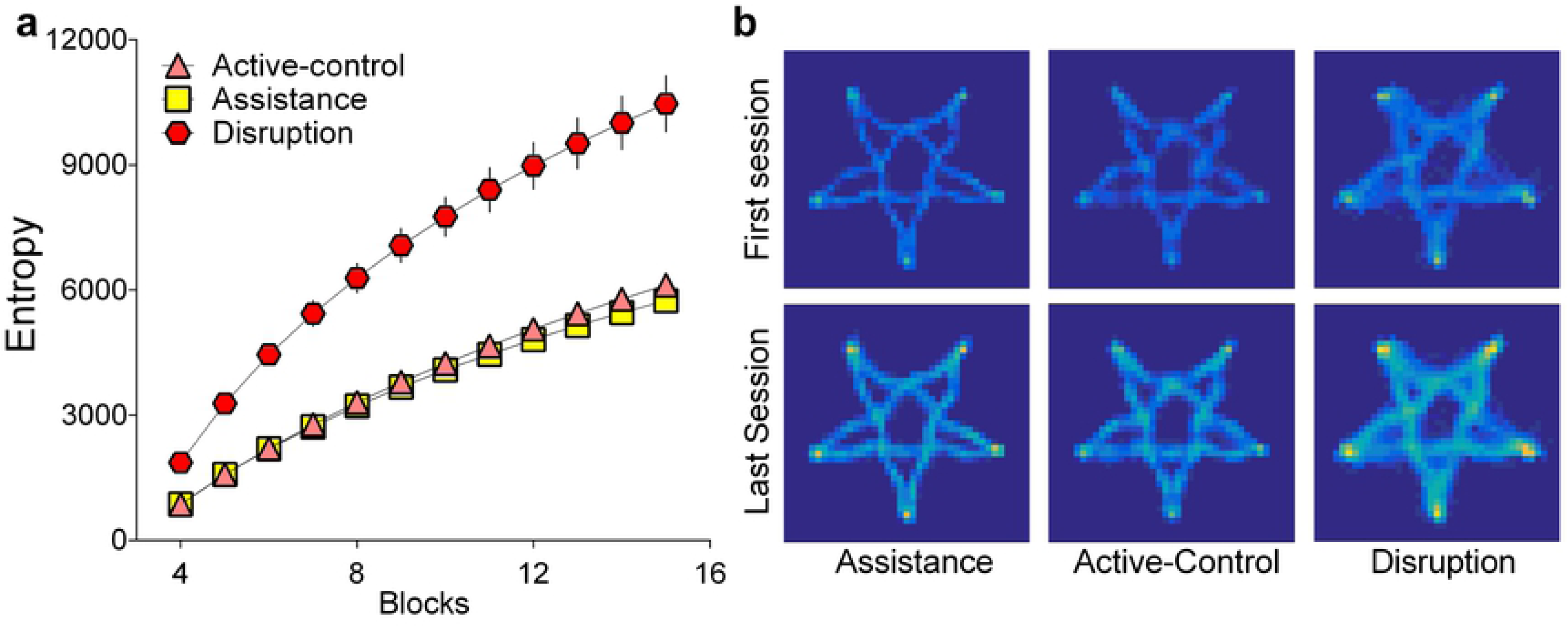
Disturbance accelerates skill acquisition. (a) Disturbance force training produced a steeper exponential performance curve during the training blocks. (b) The Disturbance training group were able to generalize their learning better than Assistance and Active Control groups, as measured by reduction in mean path error between pre- and post-tests

## Experiment 2 – Manipulating Information Sampling Without Facilitating a Short Term Impedance Strategy

The results from Experiment 1 indicate that disturbance results in faster learning in a manner consistent with the hypothesised information-driven process. However, these results do not rule out the possibility that it is disturbance forces per se that facilitate learning. For example, in Experiment 1, the adoption of a short term global impedance strategy (e.g. stiffening arm in all directions when an unexpected force was encountered) in response to disturbance forces could not be ruled out (see [25]). In Experiment 2, we therefore created algorithms that varied the amount of stiffness between trials to facilitate or constrain workspace information acquisition, and importantly make it improbable that the adoption of a global impedance strategy could yield better performance (**Fig 5A-C** and **Fig 7A-C**). The Random training condition exposed participants to an environment with a large degree of uncertainty (i.e. larger magnitude of changes in stiffness and more frequent switches between positive and negative stiffness on a trial-by-trial basis), but with an average level of overall stiffness that was close to zero. This means development of a global impedance strategy would hinder performance under the random condition (as 50% of participants’ trials were assisted with the virtual spring on average). It follows that a global impedance explanation would not account for improved performance, but the unpredictability of the stiffness between trials would induce a greater range of workspace sampling and provide the most amount of information. Thus, improved performance could be attributed to the increased exposure to information rather than the adoption of global impedance. In summary, if our hypothesis has merit then it would predict that the Random condition should lead to the best learning, whilst AA and ADA would impair learning (as they constrain information sampling).

**Fig 5.**
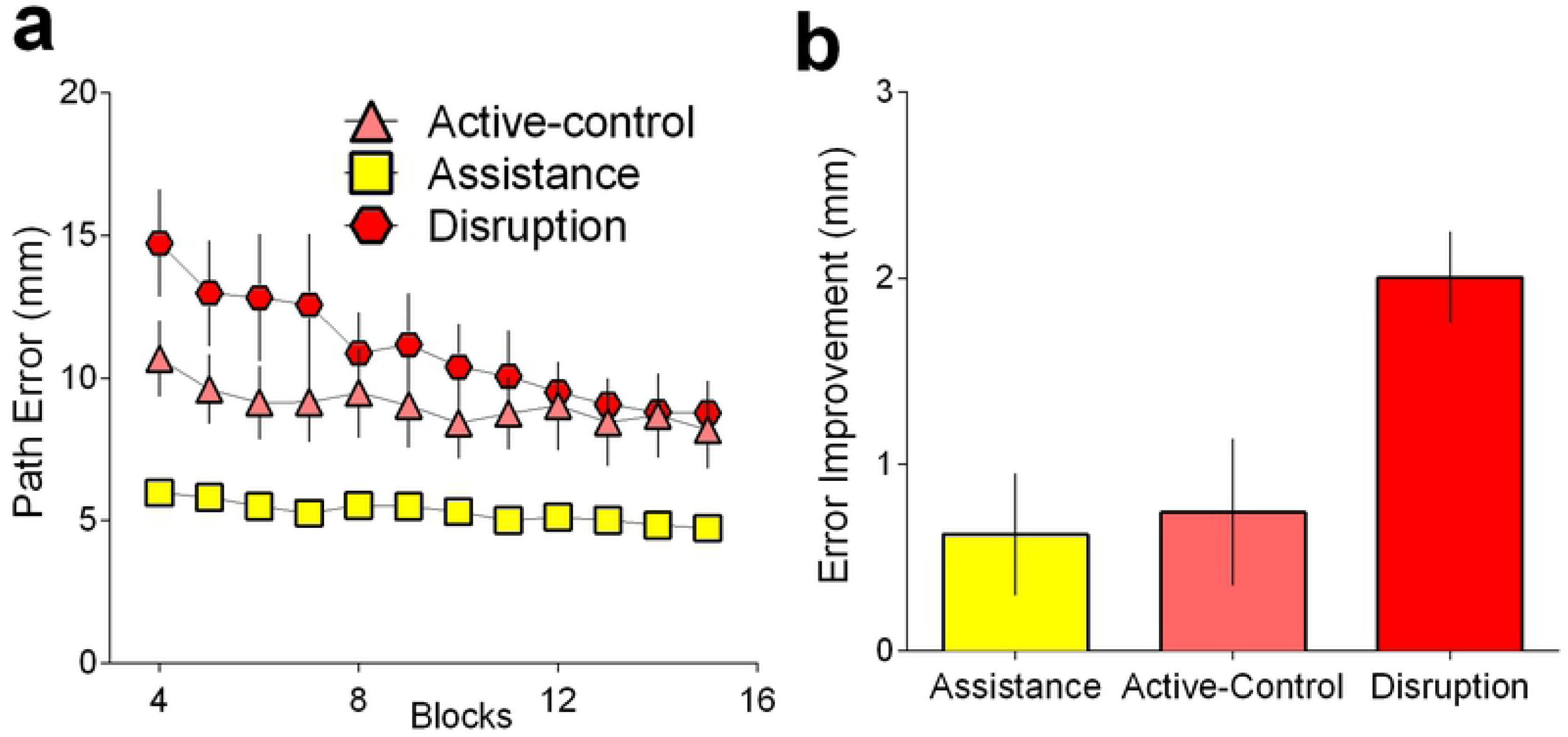
Emergent properties of the training algorithms. The level of assistance (positive stiffness) or error enhancement (negative stiffness) during training was varied on a trial-by-trial basis per the participant’s allocated group. We reasoned that the increased changes in stiffness (panel a shows magnitude of mean stiffness change between trials plotted) and switching between positive to negative stiffness values (panel b shows group average number of switches throughout training plotted) afforded to the random group would result in increased workspace information sampling and therefore greater surprise, through means other than the provision of a high negative stiffness (panel c shows average stiffness per condition).

**Fig 6.**
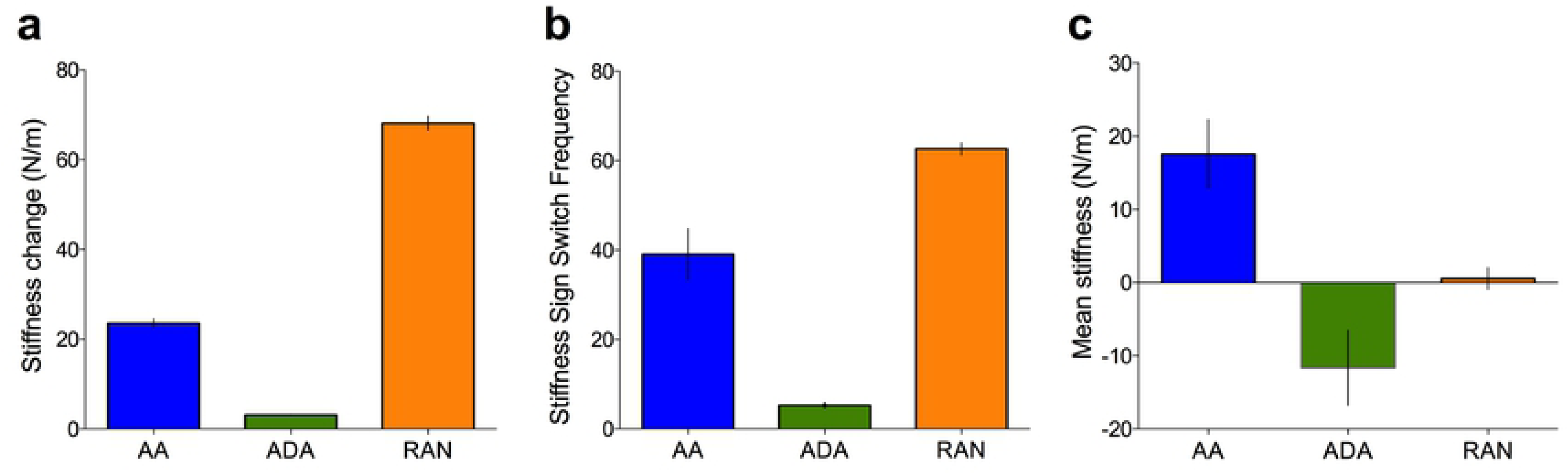
Workspace information and surprise. The stiffness coefficient *K* (N/m) demonstrates the degree of assistance (positive values/error reduction) and disturbance (negative values/error amplification) on a trial-by-trial basis for example subjects in the (a) Adaptive Algorithm; (b) Adaptive Disturbance Algorithm and (c) Random conditions; (d) The manipulation led to the Random group having more information over time; and (e) Heat maps of the amount of information across the workspace provide a visualization of difference effect for example participants, after the first and last training session.

**Fig 7.**
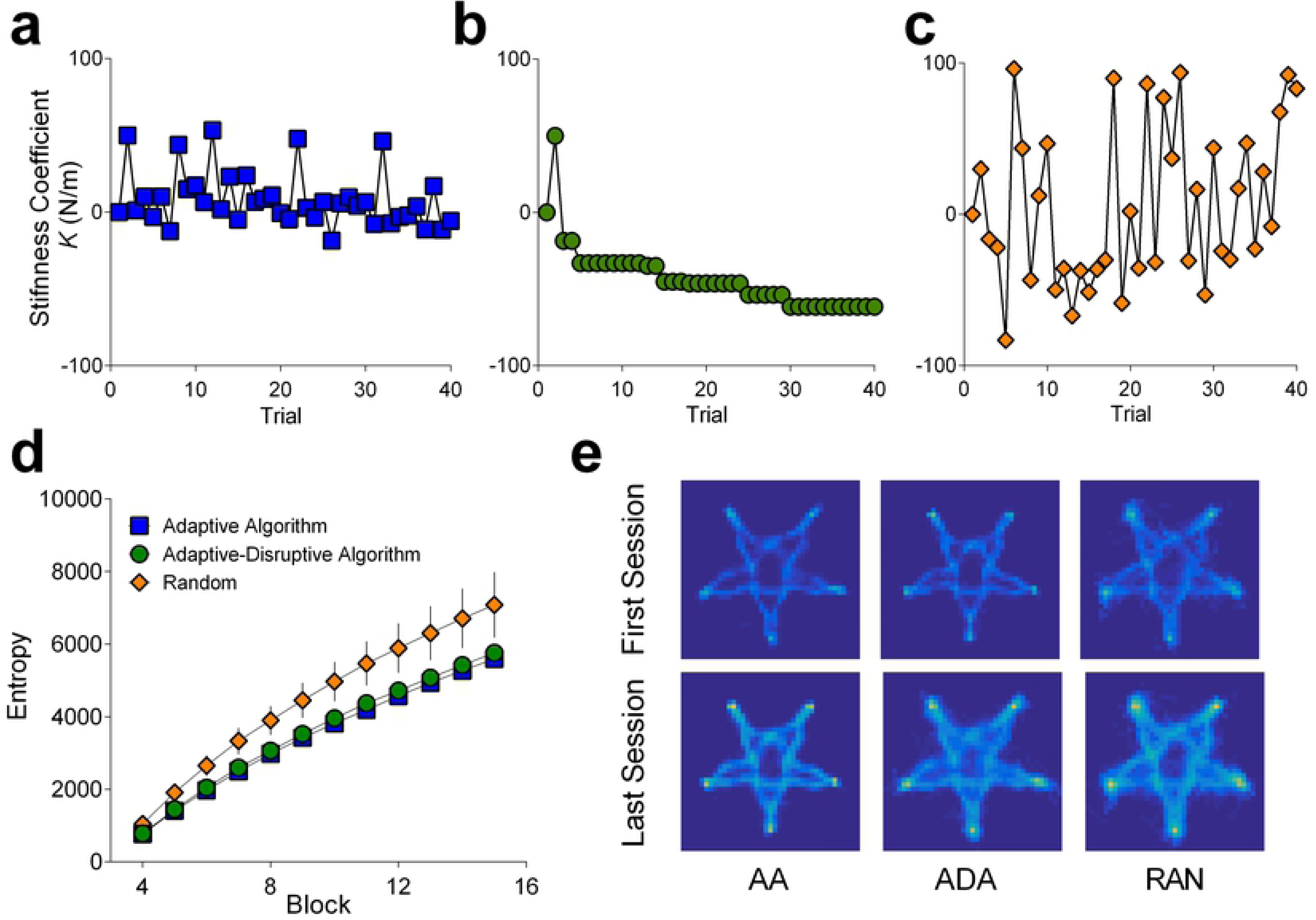
Performance on training and learning generalization. (a) Error reduction rates during training. Abscissa represents block number; (b): Random levels of assistance/disturbance demonstrated better learning, as indexed by the amount of error reduction post training relative to pre-training in a novel workspace. Pre- and post-training assessments are always performed without any stiffness intervention (k=0).

In line with our experimental aims, the algorithms produced significantly different mean values of stiffness throughout training (F (2, 41) = 12.40, p < .0001, *η*^2^_*p*_ *=* .377), mean trial-on-trial stiffness change (F (2, 41) = 931.9, p < .0001, *η*^2^_*p*_*=* .986), and number of times the task switched from assistive to disruptive (or vice versa) (F (2, 41) = 67.25, p < .0001, *η*^2^_*p*_*=* .7664).

Our predictions regarding information differences were borne out with statistically reliable group differences in the cumulative amount of workspace information at the end of training (F (2, 42) = 20.06, p < .0001, *η*^2^_*p*_*=* .489; **Fig 6D**). The Random group experienced more information relative to the Adaptive-Algorithm (p < .0001) and Adaptive-Disturbance (p < .0001) conditions, but there was no difference between the latter two groups (p = .806).

From the curve fitting results, there were no reliable differences in task difficulty level as indexed by individual values (F (2, 42) = 1.491, p = 0.2368, *η*^2^_*p*_*=*.066), but the groups did show differences in performance improvement across training (F (2, 42) = 5.058, p = .0108, *η*^2^_*p*_*=* .194). This effect was driven by the Random group showing a steeper curve in training performance compared to the Adaptive Algorithm (p = .0112), though it did not reach the statistical significance threshold when compared against the Adaptive Disturbance Algorithm (p = .0624). There were no differences between the Adaptive Algorithm and the Adaptive Disturbance conditions (p = .8613).

We also found group differences in the amount of motor learning from pre-to post-training with no stiffness intervention (F (2, 42) = 4.541, p = .0164, *η*^2^_*p*_*=* .178; **Fig 7B**). There was no statistically reliable difference in learning between the Adaptive Algorithm and Adaptive-Disturbance Algorithm (p = .914). Instead, this effect was driven by improvements following exposure to Random levels of assistance/disruption relative to the Adaptive (p = .018) and Adaptive-Disturbance algorithms (p = .009).

Finally, given our hypothesis that the amount of information predicts learning, we reasoned that there should be a positive correlation between the amount of information that participants are exposed to during training and the amount of learning (i.e. difference in performance between pre- and post-training sessions). Conducting correlation analyses at a condition-level would have been confounded by our manipulations of information across training groups and would have relatively weak statistical power to detect an underlying relationship (sample sizes varying from 13 to 17 per group). Thus, we pooled data across both experiments (n = 86) and performed a simple linear regression to predict learning based on cumulative information exposure during training. Consistent with our hypothesis, we found a statistically significant relationship (F (1, 82) = 10.45, p = .0011), with the information metric explaining 11.2% in variation in learning across all conditions (R^2^ = 0.112; **Table 1**; **Fig 8**).

**Table 1.** Information exposure predicts learning.

| Model | *t* | *p* | $\beta$ | *F* | *df* | *p* | *mult.*<br>*R*<sup>2</sup> | *adj. R*<sup>2</sup> |
| --- | --- | --- | --- | --- | --- | --- | --- | --- |
| Model 1 |  |  |  | 10.45 | 83 | 0.002 | 0.112 | 0.101 |
| Cumulative information | 3.232 | 0.002 | $2.31 \times 10^{-4}$ | | | | | |
| Model 2 |  |  |  | 1.125 | 83 | 0.292 | 0.014 | 0.001 |
| Path Error Mean SD | 1.061 | 0.292 | $6.45 \times 10^{-2}$ | | | | | |
| Model 3 |  |  |  | 5.34 | 82 | 0.006 | 0.116 | 0.094 |
| Cumulative information | 3.087 | 0.003 | $2.59 \times 10^{-4}$ | | | | | |
| Path Error Mean SD | -0.633 | 0.529 | $-4.27 \times 10^{-2}$ | | | | | |

**Fig 8.**
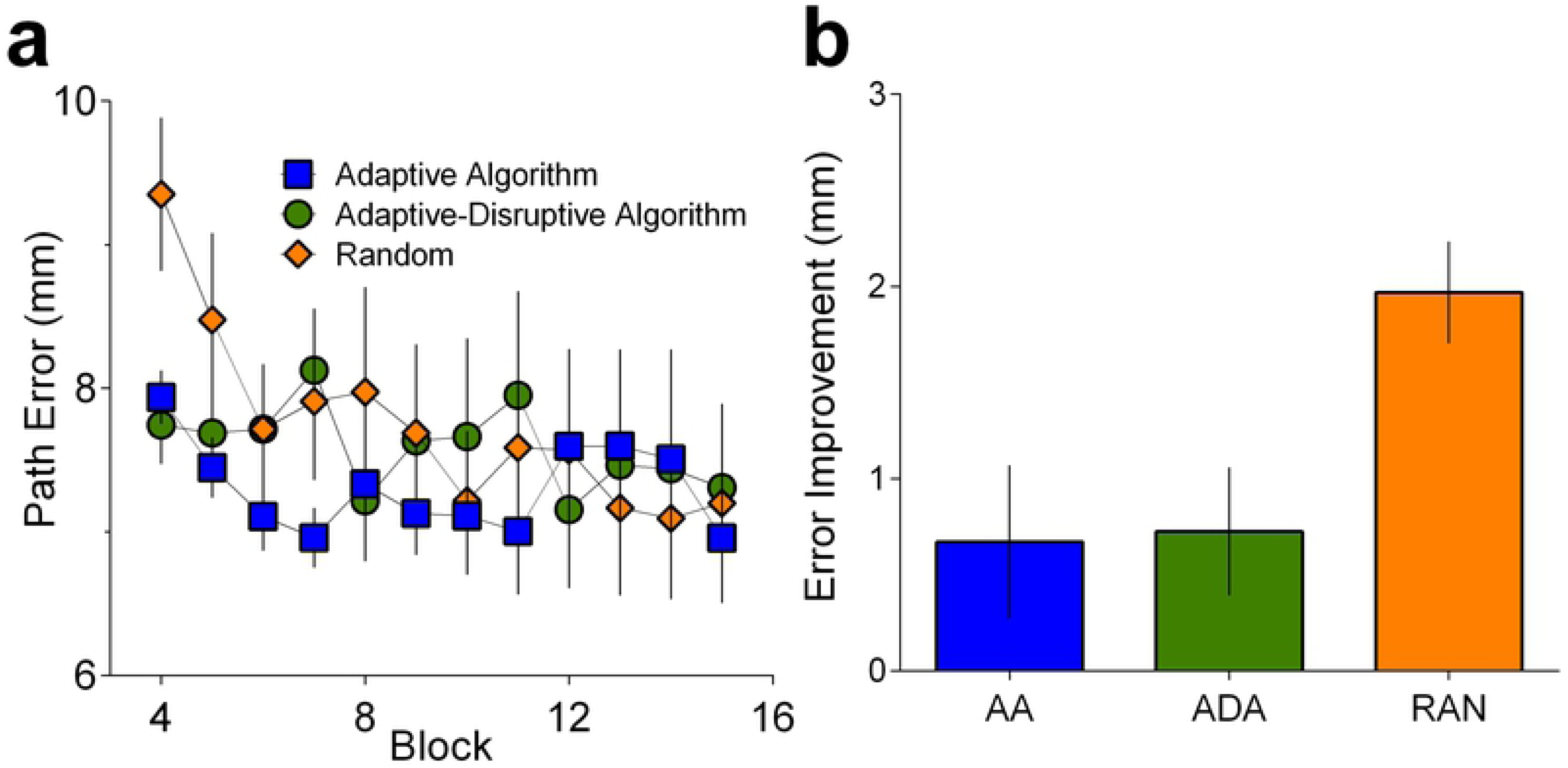
Information exposure predicts learning. Learning (mean path error reduction between pre- and post-training) as a function of cumulative information acquired during training (total entropy), for all participants in both experiments (R^2^ = 0.122).

Recent evidence from Wu and colleagues [29] demonstrates that the intrinsic movement variability associated with motor commands (from Z_n_ to Z_n+1_ to Z_n+2_ …) predicts individual rates of motor learning. Indeed, it is possible that increased error variability may be the mechanism by which information about the workspace is acquired. To contextualise and compare the predictive value of the information metric against a more parsimonious model of movement variability, we ran a second regression analysis where we included the standard deviation of path error (per component/sub-path; and averaged across training trials; **Table 1** Model 2). Interestingly, we found that this measure of variability was unable to predict learning in these data (p = .292, R^2^ = 0.01) and a direct comparison between a two-parameter model (Model 3; R^2^ = 0.116) and Model 1 showed no statistically significant reliable differences (p = .529).

## Discussion

To date, there have been no principled explanations as to why motor learning can be impaired by haptic assistance and facilitated by disturbance force application [9]. The current results support the hypothesis that the underlying mechanism relates to the availability of information, and show that haptic forces that provide more ‘surprise’ will lead to better learning in novel environments.

We created a model (inspired by the Free Energy Principle) to quantify the amount of information available to learners during a task. Experiment 1 showed that disturbance forces led to the accumulation of significantly more information across the training period. These results aligned with our analysis of the amount of motor learning following training, whereby the group that sampled more information showed superior performance relative to a group provided with assistance and to an active-control group. In Experiment 2, we demonstrated that the manipulation of information (created by training individuals on a series of random assistive and disturbance forces) yielded better learning compared to providing predictable levels of assistance/disturbance tuned to individual performance. It should be noted that the results from Experiment 2 cannot be explained by the adoption of a short term global impedance strategy (without much special pleading).

Our findings are consistent with previous results suggesting that disturbance forces might be beneficial for motor learning [4–7]. Importantly, the current work advances these reports by providing, and testing, a theoretical account of why disturbance might accelerate learning. Specifically, we show that these results are predicted by the free energy principle - which proposes that human learning can be conceptualised as a process of free-energy minimization [14]. Here, motor learning is seen as a process of entropy reduction where the average surprise of perceptual outcomes sampled from a probability distribution relating to a motor command is decreased through the development of forward models. The decrease in surprise relates to improved inferences created by the system through exposure to information that relates perceptual output to motor signal input. In line with this, through pooling the data across both experiments, we found that the amount of workspace information participants were exposed to during training could predict a statistically significant amount of variance in learning. Given the plethora of variables that could also have influenced learning across these different manipulations (six experimental conditions in two experiments), it is notable that this relationship between information and learning could be detected.

Moreover, we provide evidence that the improved information sampling created by disturbance enables generalisation rather than simple performance facilitation [1,30]. Our work thus complements and advances previous observations about the potential benefits of disturbance. For example, an earlier study showed that performance on a tracking task could be improved through delivery of haptic disturbance [5]. This finding could be explained, however, by the participants being trained to become more proficient in deploying feedback control and, indeed, the authors of the study explained their results in terms of a general training improvement in the ‘attentional’ capabilities of their participants. The problem with such explanations relates to the difficulty in defining and quantifying the term ‘attention’ when used in this manner. It is therefore interesting to note that the improved tracking performance is predicted within the FEP framework. The presence of haptic disturbance when tracking will generate surprise and thus force the system to act to reduce the entropy (i.e. learn to make effective feedback corrections). Indeed, the Random training condition in our experiment exploited this mechanism in a principled manner by exposing participants to frequent movement-by-movement switches between positive and negative stiffness. Together, these results illustrate the fundamental links between attention and uncertainty (see [31,32]), and suggest that the effects of haptic disturbance can be quantified in a range of different settings through information theory.

Our results also build on previous work showing a relationship between variability and motor learning. For example, Van Beers [33] showed that the random effects of planning noise accumulate, in contrast to task-relevant errors which show close to zero accumulation (explained by effective trial-by-trial corrections), whilst Wu et al’s experiments [29] (results described earlier), have shown that task-relevant motor variability facilitates faster learning rates. On these grounds, it has been argued that intrinsic movement variability leads to motor exploration, which sub-serves motor learning and performance optimization. Indeed, the idea that action exploration can drive learning has long been mooted in theories of operant behaviour [34] and human development [35–37]. Recent experiments have shown that (a) artificially manipulating the relationship between movements and visuomotor noise can be used to teach people specific control policies [38] and (b) the variability in task-redundant parameters can predict motor adaptation rates [39]. The current findings demonstrate that extrinsic variability delivered through haptic disturbance can, in the same vein, augment learning by increasing the amount of information sampled by the learner. The general notion that increased exposure to information can lead to faster learning is well explained by theories of structural learning and has good support from a range of empirical studies [18,19,40–43] including investigations of laparoscopic surgical training [44]. Our extension to these ideas is that learning of the structure can be directly related to the amount of information available to the learner. Indeed, regression analyses for our data shows that the amount of information accumulated over training (as indexed by our model) provided greater explanatory power compared to a measure of motor variability alone in this task.

These findings raise the issue of which neural substrates underpin these learning processes. The neural processes that implement the computational algorithms exploited by the human nervous system remain to be discovered [45,46]. Likewise, the underlying control mechanisms supporting skilled arm movements are poorly understood and, as such, it is difficult to speculate on how the individuals learned to compensate for the complex force field, but we suggest that the learning was likely to involve processes related to optimal feedback control as well as predictive mechanisms [47–49].

Our findings suggest that the participants developed forward or inverse models that allowed them to predict (and thus compensate for) the novel force field through which they needed to move. It has been shown previously that participants can learn a short term strategy of stiffening their arm to resist the effects of sudden unexpected force perturbations [24,25]. This work has demonstrated that humans learn to use selective control of impedance geometry in order to stabilise unstable dynamics in a skilful and energy efficient manner. It is probable that participants in the current experiments adopted such a strategy when they were first exposed to the novel workspace (as they were unable to predict the forces that were applied as they moved through the space). Importantly, there was a regular (lawful) structure to the novel workspace, in the same way that the world provides a lawful force field through which the neonate must learn to move their arm. We hypothesised that the system would learn the underlying force field so that the arm could move skilfully through the workspace rather than repeatedly contend with unexpected displacement. This hypothesis was based on the free energy minimization principle which suggests human behaviour is marked by continual attempts to reduce entropy (i.e. minimise surprise). Experiment 2 allowed us to test whether participants were learning the force field or adopting a global impedance strategy, by which the arm is stiffened in all directions to counteract external force interventions. As outlined above and demonstrated in previous research, participants are likely to adopt a global impedance strategy when the force intervention is largely disruptive and increases error (k < 0). However, in Experiment 2, the random condition consisted of (on average) 50% assistive trials, whereby the force intervention *assisted* movement, thus rendering such a strategy sub-optimal. We reasoned that, in contrast to the random forces, the adaptive disturbance algorithm, where participants were provided with a more consistent presentation of disturbance forces would be more likely to adopt an impedance control strategy. Given that we observed improved learning in the random condition, impedance control is unlikely to provide a full account of these data. Instead, these results indicate that participants were learning to skilfully counteract the underlying workspace force field and we propose that this learning was promoted, in part, through the increased information acquired during training.

Finally, it is important to note that this study used neurologically intact adults as participants and whilst the force field in the two experiments allowed us to examine novel skill learning, the difficulty was tuned to a level such that all participants could complete the task. We speculate that disrupting the training of individuals with neurological deficits (e.g. cerebral palsy) might not be beneficial, and constraining errors in these populations could speed up learning by helping the individuals sample the necessary information [21]. Consistent with this, there is work with stroke survivors that has shown that error amplification is useful in rehabilitation for mild impairment, but error guidance is necessary for patients with more severe damage [50]. Likewise, haptic guidance has been found to be beneficial for people with relatively low skill levels, but error enhancement is better for highly skilled individuals [3,51]. The current work builds on these observations and provides a theoretical framework for the development of optimized robotic training devices in skill training and rehabilitation.

## Author Contributions

E.J. & J.B. conducted the observations. E.J. and J.B. reduced the data. F.M., J.B, and M.M-W wrote the manuscript. All authors discussed the results and implications and commented on the manuscript at all stages.

## Data Availability

Data files used in analyses are openly available on the Open Science Framework: https://osf.io/7c95b/

## Acknowledgements

We would like to thank our participants for taking part in the reported experiments. F.M and M.M-W were supported by Fellowships from the Alan Turing Institute and a Research Grant from the EPSRC (EP/R031193/1).

